# PIKfyve inhibition induces LRRK2 activation through phosphatidylinositol 3-phosphate and CASM

**DOI:** 10.64898/2025.12.16.694573

**Authors:** Tomoki Kuwahara, Gen Yoshii, Maria Sakurai, Shoichi Suenaga, Hiroki Nakanishi, Matthew Jefferson, Thomas Wileman, Kentaro Tomii, Takeshi Iwatsubo

**Author notes:** Corresponding authors at: 7-3-1, Hongo, Bunkyo-ku, Tokyo, 113-0033, Japan. These authors contributed equally to this work (Tomoki Kuwahara, Gen Yoshii).

## Abstract

LRRK2 is a Parkinson’s disease (PD)-associated kinase that phosphorylates Rab GTPases. Hyperactivation of LRRK2, which is thought to cause PD, occurs on stressed endolysosomal membranes via CASM (conjugation of ATG8 to single membranes) and Rab, whereas the involvement of specific membrane lipids remained unclear. Here, we found that the inhibition of PIKfyve that metabolizes phosphatidylinositol 3-phosphate (PI3P) induces hyperactivation of LRRK2, and that additional inhibition of the class III phosphoinositide 3-kinase (PI3K) that generates PI3P suppressed LRRK2 hyperactivation. Cellular PI3P levels were indeed elevated when LRRK2 was activated. CASM-induced activation of LRRK2, but not CASM itself, was suppressed by PI3K inhibitors, whereas inhibition of CASM or Rab12/Rab29 suppressed LRRK2 activation induced by PIKfyve inhibitors. LRRK2 and phospho-Rab10 were co-localized with PI3P on LAMP1-positive enlarged vacuoles, and LRRK2 directly bound PI3P in vitro via its N-terminal positively charged amino acids. Together, we propose an updated mechanism of LRRK2 hyperactivation that simultaneously requires CASM, Rab and PI3P-rich membranes.

## Introduction

Leucine rich repeat kinase 2 (LRRK2) is a large serine/threonine kinase that phosphorylates a subset of Rab GTPases in cells (Alessi and Pfeffer, 2024). LRRK2 consists of 2527 amino acids and harbors multiple functional domains, including the kinase domain, the Roc (Ras of complex proteins)-COR (C-terminal of Roc) domain, the LRR domain, the N-terminal Armadillo (ARM) and Ankyrin (ANK) repeats, and the C-terminal WD40 domain (Iannotta and Greggio, 2021; Komori and Kuwahara, 2023). Missense mutations in *LRRK2* have been identified as a major cause for autosomal-dominant Parkinson’s disease (PD) (Paisán-Ruíz et al., 2004; Zimprich et al., 2004). The well-established familial mutations reported to date are all located either in the kinase domain (G2019S, I2020T) or the Roc-COR domain (N1437H, R1441C/G/H, Y1699C), and structural analyses using cryo-electron microscopy have revealed the close proximity of these two domains (Myasnikov et al., 2021; Zhang and Kortholt, 2023). All these pathogenic mutations have been shown to significantly enhance the Rab phosphorylation activity of LRRK2 (Steger et al., 2016), leading to the hypothesis that abnormal hyperactivation of LRRK2 causes PD.

LRRK2 is widely expressed in the body, with particularly high expression in the brain, lung, kidney, and immune cells such as macrophages and neutrophils (Fan et al., 2018; Gardet et al., 2010; Giasson et al., 2006). In the brain, LRRK2 is expressed ubiquitously in neurons, astrocytes and microglia. Among these cell types, macrophage lineage cells express high levels of LRRK2 upon exposure to inflammatory stimuli, especially interferon-γ (IFN-γ) (Gardet et al., 2010; Thévenet et al., 2011). Genetic variants in *LRRK2* have also been associated with Crohn’s disease and leprosy (Barrett et al., 2008; Zhang et al., 2009). These reports point to the potential importance of LRRK2 in immune-related cells, particularly in macrophage lineage.

Research on LRRK2 has been extensively conducted in the PD research field, and in clinical settings where LRRK2 inhibitors have been explored intensively (Jennings et al., 2023). In basic biological research, the focus has been on the mechanism of LRRK2 hyperactivation associated with PD and the downstream mechanisms caused by the enhanced Rab phosphorylation. It has been reported that excessive Rab phosphorylation leads to lysosomal dysfunctions indicated by altered morphology or positioning, as well as ciliary defects caused by abnormal centrosome function (Fasiczka et al., 2023; Kuwahara and Iwatsubo, 2020). We and others have shown that LRRK2 is activated on lysosome membranes when lysosomes are stressed by chloroquine (CQ), monensin, nigericin or L-Leucyl-L-Leucine methyl ester (LLOMe) (Bonet-Ponce et al., 2020; Eguchi et al., 2018; Herbst et al., 2020; Kalogeropulou et al., 2020). This activation can be mediated by several Rab GTPases, such as Rab32 family members (Rab29, Rab32, Rab38) and Rab12 (Dhekne et al., 2023; Gustavsson et al., 2024; Hop et al., 2024; Purlyte et al., 2018; Unapanta et al., 2023; Wang et al., 2023a). LRRK2 is also activated during conjugation of ATG8 to single membranes (CASM), which is induced by pH elevation and ionic/osmotic imbalances within lysosomes (Florey et al., 2015; Jacquin et al., 2017) and is primarily mediated by the V-ATPase-ATG16L1 axis (Durgan and Florey, 2022; Wang et al., 2022). CASM mediates activation by recruiting LRRK2 and ATG8 family proteins onto the endolysosomal membranes (Eguchi et al., 2024; Kuwahara and Iwatsubo, 2024), and this may involve binding of GABARAP to LRRK2 (Bentley-DeSousa et al., 2025b). In both mechanisms, Rab and GABARAP have been shown to bind to specific sites within the N-terminal ARM repeats of LRRK2, thereby anchoring LRRK2 to the endolysosomal membranes (Bentley-DeSousa et al., 2025a; Sakurai and Kuwahara, 2025).

LRRK2 has also been shown to bind directly to negatively charged lipids in membranes (Wang et al., 2023b). Since LRRK2 becomes activated on the membranes of stressed lysosomes, specific lipids may also contribute to LRRK2 activation, although such a mechanism has remained unexplored. Endolysosomes mainly contain two phosphoinositide lipids: phosphatidylinositol 3-phosphate (PI3P) and phosphatidylinositol 3,5-bisphosphate (PI(3,5)P_2_), with the ratio of PI(3,5)P_2_ increasing as endosomes mature (Posor et al., 2022; Wallroth and Haucke, 2018). We found that treating cells with PIKfyve inhibitors that block PI3P metabolism to PI(3,5)P_2_ potently activate LRRK2, leading us to study the relationship between phosphoinositide changes and LRRK2 activation. The results offer a novel activation model where PI3P is involved in LRRK2 activation in addition to CASM and Rab.

## Results

### LRRK2 is activated in response to pharmacological stimuli that cause an increase in PI3P

To explore the endolysosomal lipid component that influences LRRK2 kinase activity, we focused on PIKfyve, a lipid kinase that regulates endolysosomal function by generating PI(3,5)P_2_ from PI3P (Rivero-Ríos and Weisman, 2022). Inhibition of PIKfyve has been shown to induce the formation of swollen endolysosomal vacuoles in cells (Krishna et al., 2016; Uwada et al., 2025), which resemble changes induced by treatment with chloroquine (CQ) or monensin, known pharmacological activators of LRRK2. We used mouse macrophage RAW264.7 cells pretreated with IFN-γ, as this treatment greatly increases the expression of endogenous LRRK2 in these cells (Gardet et al., 2010), enabling sensitive detection of LRRK2 activity changes (Eguchi et al., 2018; Kuwahara et al., 2020). Incubation of these cells with apilimod or YM201636, two widely used inhibitors of PIKfyve (**Fig. 1A**), for 2 hours caused the formation of swollen endolysosomal vacuoles positive for LAMP1 (**Fig. 1B**). We found that these treatments also caused marked increase in the kinase activity of LRRK2, as measured by the phosphorylation of its substrate Rab GTPases including Rab10 (**Fig. 1C and D**). This phosphorylation is dependent solely on LRRK2, as treatment with an LRRK2 inhibitor MLi-2 abolished Rab10 phosphorylation **(Supplementary Fig. 1A)**. As an orthogonal approach, we treated cells with two different small interfering RNAs (siRNAs) specific to PIKfyve and confirmed the increase in Rab10 phosphorylation as well as PIKfyve knockdown after 48 hours of treatment **(Supplementary Fig. 1B and C**). To determine whether this LRRK2 hyperactivation resulted from a decrease in PI(3,5)P_2_ or an increase in PI3P due to PIKfyve inhibition, we treated cells with inhibitors of class III phosphoinositide 3-kinase (PI3K) that generates PI3P from phosphatidylinositol (PI) (**Fig. 1A**). The results showed that the elevation of LRRK2 activity upon PIKfyve inhibition was suppressed by treatment with a selective class III PI3K inhibitor VPS34-IN1 (**Fig. 1C and D**), a second class III PI3K inhibitor Compound 19 (**Fig. 1E and F**), or by another PI3K inhibitor 3-methyladenine (3-MA) **(Supplementary Fig. 1D and E**). These results suggest that an increase in PI3P, rather than a decrease in PI(3,5)P_2_, is involved in LRRK2 hyperactivation. As a note, treatment with class III PI3K inhibitors alone, especially VPS34-IN1, in the absence of PIKfyve inhibitors did not reduce the low basal level of Rab10 phosphorylation but rather increased it (*P* = 0.0025 for VPS34-IN1) (**Fig. 1C-F**), implying that low LRRK2 activity under steady-state conditions is also influenced in a complex manner by factors other than PI3K.

**Figure 1.**
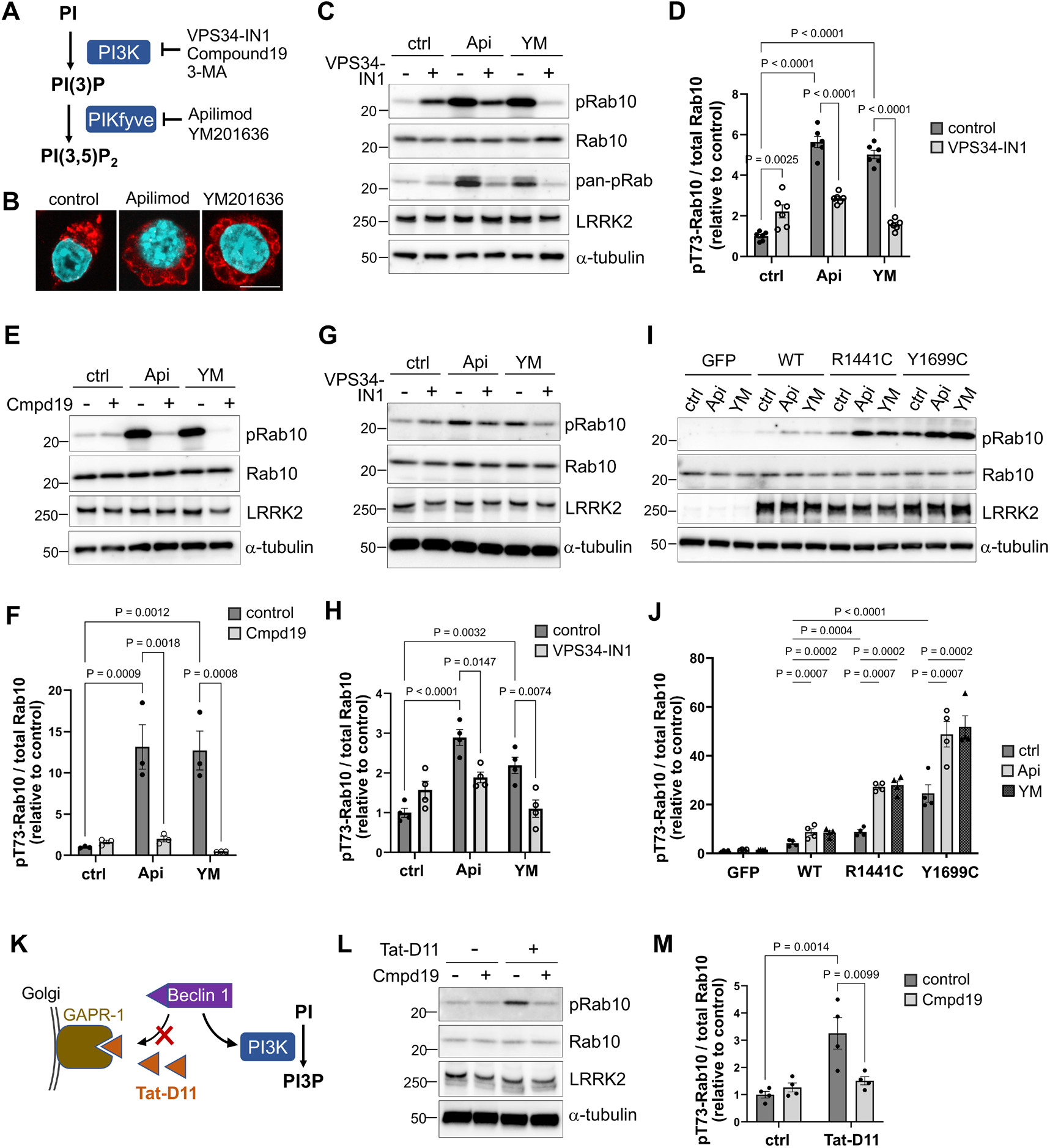
LRRK2 is activated in response to pharmacological stimuli that cause an increase in PI3P. **(A)** Schematic diagram of the PI3P metabolic pathway, where class III PI3K is responsible for the generation of PI3P from PI and PIKfyve is for the generation of PI(3,5)P_2_ from PI3P. Compounds that inhibit each kinase are also shown. **(B)** Immunostaining of intracellular vacuoles formed by treatment with apilimod (Api) or YM201636 (YM) in RAW264.7 cells. Vacuoles were stained with anti-LAMP1 antibody (red) and nuclei were stained with DAPI (cyan). Scale bar = 10 μm. **(C-F)** Immunoblot analysis of Thr73-phosphorylated Rab10 (pRab10) and other indicated proteins in RAW264.7 cells treated with PIKfyve inhibitors as well as VPS34-IN1 (C, D) or Compound19 (Cmpd19) (E, F). Bar graphs in D and F show quantification of pRab10 relative to total Rab10. Data represent mean ± SEM, n = 6 (D) or 3 (F), and the difference was analyzed using two-way ANOVA with Tukey’s test. **(G, H)** Immunoblot analysis of pRab10 and other proteins in A549 cells treated with PIKfyve inhibitors as well as VPS34-IN1. A bar graph in H shows quantification of pRab10 relative to total Rab10. Data represent mean ± SEM, n = 4, two-way ANOVA with Tukey’s test. **(I, J)** Immunoblot analysis of pRab10 in HEK293 cells that overexpress GFP, LRRK2-wild type (WT) or pathogenic mutants (R1441C, Y1699C) and are treated with PIKfyve inhibitors. A bar graph in J shows quantification of pRab10/total Rab10. Data represent mean ± SEM, n = 4, two-way ANOVA with Tukey’s test. **(K)** Schematic illustration of the activation induction mechanism of PI3K by Tat-D11. **(L, M)** Immunoblot analysis of pRab10 and other indicated proteins upon treatment with Tat-D11 and/or Cmpd19 in RAW264.7 cells. A bar graph in M shows quantification of pRab10 relative to total Rab10. Data represent mean ± SEM, n = 4, two-way ANOVA with Tukey’s test.

Next, we examined whether similar effects could be observed in cells other than RAW264.7 cells. Human lung epithelial A549 cells are known to express high level of endogenous LRRK2 (Kamikawaji et al., 2013; Lobbestael et al., 2013). Treatment of A549 cells with PIKfyve inhibitors activated LRRK2, and this activation was suppressed by PI3K inhibition (**Fig. 1G and H**). Human embryonic kidney 293 (HEK293) cells express lower levels of endogenous LRRK2, while these cells transfected with 3×FLAG-tagged human LRRK2 exhibited elevation of LRRK2 activity upon PIKfyve inhibition, which was again suppressed by PI3K inhibition (**Fig. 1I and J**). More importantly, HEK293 cells expressing PD-associated mutant LRRK2 (R1441C, Y1699C) exhibited higher basal LRRK2 activity as well as further elevated activity upon PIKfyve inhibition, as compared to cells expressing wild-type LRRK2 (**Fig. 1I and J**). These results suggest that LRRK2 activation by PIKfyve inhibition occurs regardless of cell type and may represent a phenotype potentially relevant to PD.

We further examined whether the upregulation of PI3K activity to enhance PI3P generation is sufficient to induce LRRK2 activation. Tat-D11 is an improved form of the class III PI3K-activating peptide Tat-Beclin 1, which facilitates the incorporation of Beclin 1 into the PI3K complex by interfering with its binding to Golgi-localized GAPR-1 (Chang et al., 2019; He et al., 2022; Zhao et al., 2021) (**Fig. 1K**). We found that the Tat-D11 treatment in RAW264.7 cells induced the upregulation of Rab10 phosphorylation as well as LC3 lipidation (**Fig. 1L and M, Supplementary Fig. 1F-H)**. This upregulation of Rab10 phosphorylation was canceled by PI3K inhibition (**Fig. 1L and M**), confirming the involvement of PI3K activity in LRRK2 activation. Also, treatment with a LRRK2 inhibitor MLi-2 suppressed the upregulation of Rab10 phosphorylation but not LC3 lipidation **(Supplementary Fig. 1F-H)**, suggesting that LC3 lipidation occurs independently of LRRK2 activation under enhancement of PI3P generation.

### Hyperactivation of LRRK2 by diverse endolysosomal stimuli is accompanied by increased level of PI3P

LRRK2 is well known to be activated on the membranes of lysosomes stressed by lysosomotropic agents, ionophores or lysosome-permeabilizing agents (Bonet-Ponce et al., 2020; Eguchi et al., 2018; Herbst et al., 2020; Kuwahara et al., 2020). We then examined the possibility that LRRK2 activation in response to lysosomal stress may involve an increase in PI3P. Chloroquine (CQ) is a representative lysosomotropic agent that accumulates within lysosomes and causes their osmotic swelling (Florey et al., 2015; Marceau et al., 2012). Treatment of RAW264.7 cells with CQ activated LRRK2, and this activation was suppressed by treatment with the PI3K inhibitors VPS34-IN1 or Compound 19 (**Fig. 2A-C**). CQ treatment also induced LC3 lipidation via CASM (conjugation of ATG8 to single membranes), but this was not suppressed by PI3K inhibition (**Fig. 2A, D and E)**, consistent with previous findings that PI3K activity is not required for CASM (Fletcher et al., 2018; Florey et al., 2015).

**Figure 2.**
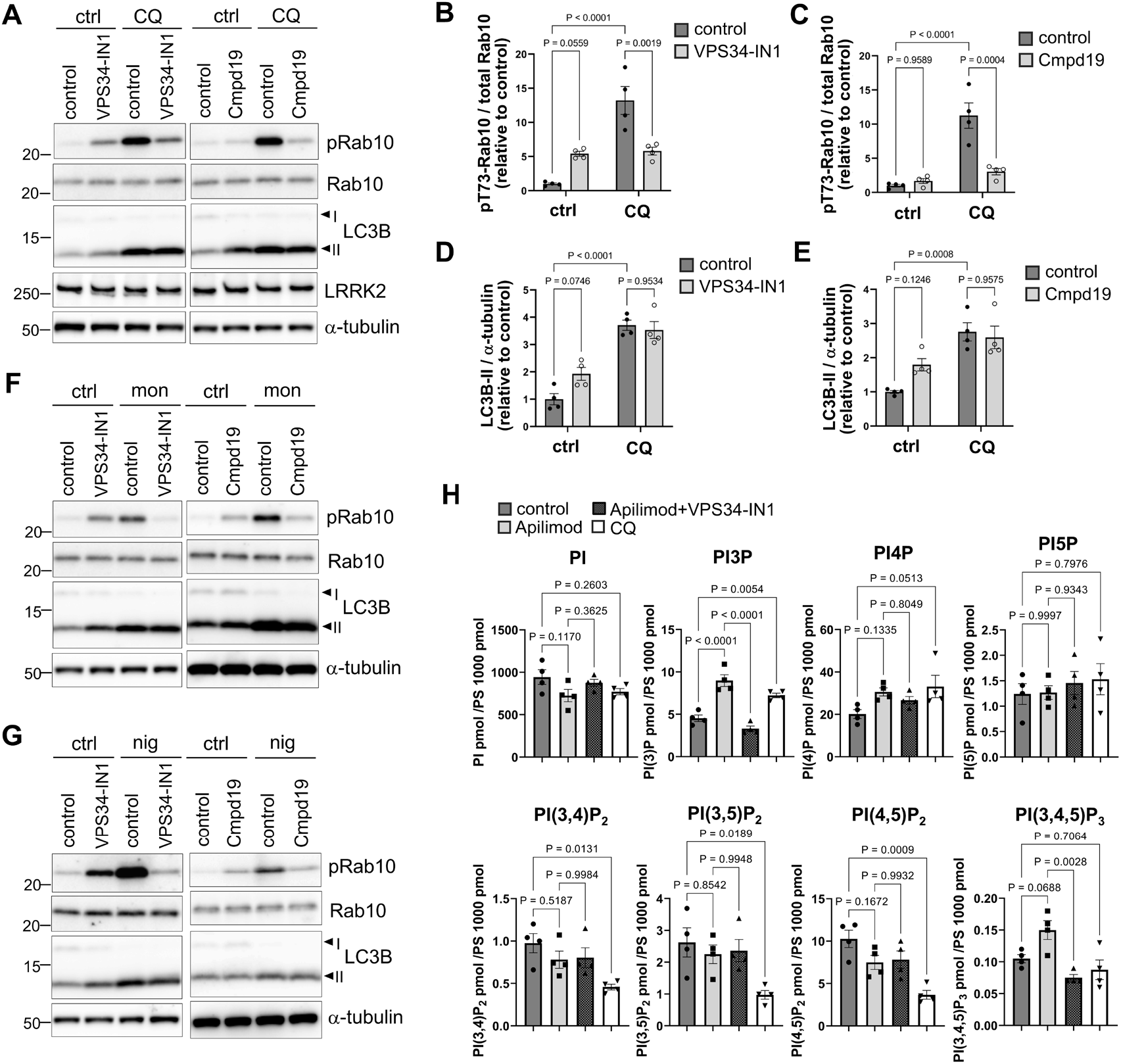
Hyperactivation of LRRK2 by diverse stimuli is accompanied by increased level of PI3P. **(A)** Immunoblot analysis of pRab10, LC3B and other indicated proteins in RAW264.7 cells treated with CQ as well as PI3K inhibitors. **(B-E)** Quantification of pRab10 relative to total Rab10 (B-C) or LC3B-II relative to α-tubulin (D-E), as shown in A, under treatment with or without CQ plus VPS34-IN1 (B, D) or Compound 19 (C, E). Data represent mean ± SEM, n = 4, and the difference was analyzed using two-way ANOVA with Tukey’s test. **(F, G)** Immunoblot analysis of pRab10 and LC3B in RAW264.7 cells treated with monensin (mon) (F) or nigericin (nig) (G) as well as PI3K inhibitors. Representative images from three independent experiments are shown. **(H)** Mass spectrometry (MS)-based measurements of phosphoinositide isomer levels in RAW264.7 cells treated with apilimod, apilimod plus VPS34-IN1, or CQ. Data represent mean ± SEM, n = 4, and the difference was analyzed using one-way ANOVA with Tukey’s test.

We next tested the effects of monensin and nigericin also known to activate LRRK2 (Kalogeropulou et al., 2020; Kuwahara et al., 2020). Both reagents act on the lysosomal membrane as monovalent cation/proton ionophores, inducing ionic and osmotic imbalances (Jacquin et al., 2017; Mollenhauer et al., 1990; Rangasamy et al., 2018). Treatment with these reagents indeed activated LRRK2, and this activation was again suppressed by the PI3K inhibitors (**Fig. 2F and G**). LC3 lipidation induced by monensin or nigericin was not suppressed by PI3K inhibition, as observed under CQ treatment (**Fig. 2F and G**). These findings suggest that lysosomal stress-induced activation of LRRK2 requires PI3P generation, while the concurrent LC3 lipidation via CASM occurs independently of PI3P generation.

Although the results above supported a link between increased PI3P levels and LRRK2 activity in cells, inhibition of PIKfyve has been shown to primarily cause a reduction in cellular PI(3,5)P_2_ levels (Cai et al., 2013; Jefferies et al., 2008). However, several studies have also shown a significant increase in PI3P levels at the same time (Cai et al., 2013; Shimanaka et al., 2022), and given that phosphoinositide (PIP) levels are regulated by a complex balance between kinase and phosphatase activities (Posor et al., 2022; Rivero-Ríos and Weisman, 2022), their changes in levels are expected to vary depending on cellular conditions and environment. Thus, we sought to perform a mass spectrometry-based quantification in our cells to confirm changes in PIP isomer levels in response to PIKfyve inhibition, PIKfyve plus PI3K inhibition, or lysosome stress. Because simple mass spectrometry cannot measure the levels of PI3P and PI(3,5)P_2_ while distinguishing them from regioisomers having the same mass (i.e., PI monophosphates (PI4P, PI5P) and diphosphates (PI(3,4)P_2_, PI(4,5)P_2_)), we employed a recently developed method that integrates the separation of regioisomers using specialized chiral chromatography (Morioka et al., 2022). As a result, PI3P levels increased upon apilimod treatment, decreased upon both apilimod and VPS34-IN1 treatment, and increased upon CQ treatment (**Fig. 2H, Supplementary Table 1)**. The levels of PI(3,5)P_2_ did not change significantly in our cells under these conditions, and the levels of other PIPs remained unchanged as well (**Fig. 2H, Supplementary Table 1)**. CQ treatment additionally caused a decrease in three phosphatidylinositol diphosphates (PI(3,4)P_2_, PI(3,5)P_2_, PI(4.5)P_2_) (**Fig. 2H, Supplementary Table 1)**. These data strengthen the notion that hyperactivation of LRRK2 involves an increase in intracellular PI3P levels.

### CASM is involved in LRRK2 activation upon PIKfyve inhibition

Next, we investigated the relationship between LRRK2 activation induced by PIKfyve inhibition and CASM, which also mediates LRRK2 activation in response to lysosomal stress (Bentley-DeSousa et al., 2025b; Eguchi et al., 2024; Kuwahara and Iwatsubo, 2024). The key factors in CASM are WD40 domain of ATG16L1, which is not required for macroautophagy but is required for binding ATG16L1 and the ATG5–ATG12 LC3 conjugation machinery to the lysosomal V-ATPase in response to raised lysosomal pH (**Fig. 3A**). Unlike macroautophagy, CASM does not require class III PI3K activity (Fletcher et al., 2018; Florey et al., 2015), and therefore it remained unclear whether CASM is involved in LRRK2 activation upon PI3P elevation.

**Figure 3.**
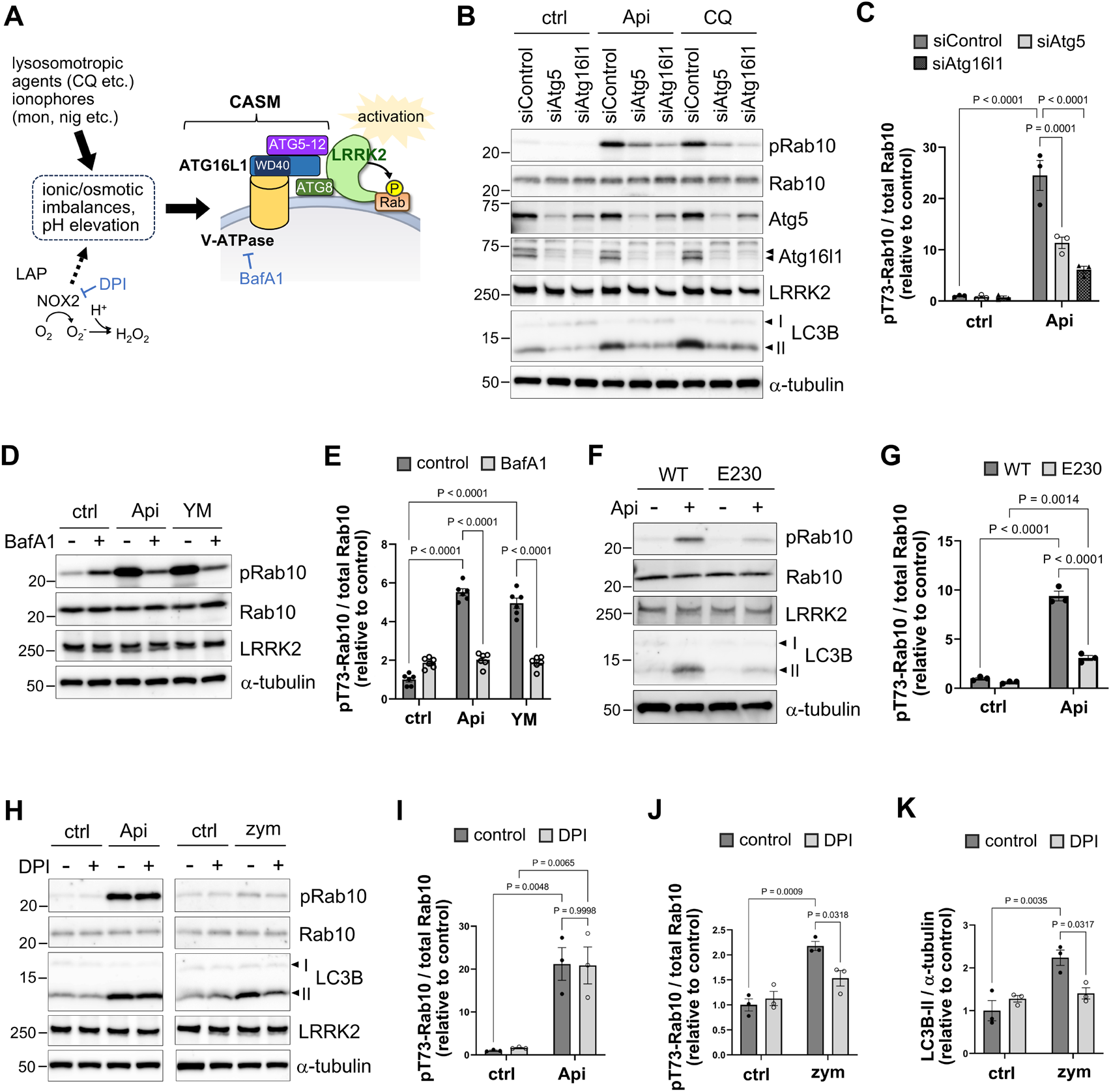
CASM is involved in LRRK2 activation upon PIKfyve inhibition. **(A)** Schematic diagram of the mechanism of the CASM-LRRK2 pathway induced by pH/ionic imbalances. LAP: LC3-associated phagocytosis. **(B, C)** Immunoblot analysis of pRab10 and other indicated proteins in RAW264.7 cells that were pretreated with siRNAs for Atg5 or Atg16l1 and then treated with apilimod or CQ. Knockdown efficiencies of Atg5 and Atg16l1 are also shown. A bar graph in C shows quantification of pRab10 relative to total Rab10, as shown in B. Data represent mean ± SEM, n = 3, two-way ANOVA with Tukey’s test. **(D, E)** Immunoblot analysis of pRab10 and other proteins in RAW264.7 cells treated with PIKfyve inhibitors as well as bafilomycin A_1_ (BafA1). A bar graph in E shows quantification of pRab10/total Rab10, as shown in D. Data represent mean ± SEM, n = 6, two-way ANOVA with Tukey’s test. **(F, G)** Immunoblot analysis of Rab10 and other indicated proteins in WT or E230 bone marrow-derived macrophages (BMDMs) treated with or without apilimod. A bar graph in G shows quantification of pRab10/total Rab10, as shown in F. Data represent mean ± SEM, n = 3, two-way ANOVA with Tukey’s test. **(H)** Immunoblot analysis of pRab10 and LC3B in RAW264.7 cells treated with apilimod or zymosan (zym) in addition to DPI. Bar graphs show quantifications of pRab10/total Rab10 upon treatment with apilimod (I) or zymosan J) and that of LC3B-II relative to α-tubulin upon treatment with zymosan (K). Data represent mean ± SEM, n = 3, two-way ANOVA with Tukey’s test.

We first examined the effect of knockdown of Atg16l1 or Atg5, both of which participate in CASM execution, on PIKfyve inhibition-induced activation of LRRK2. Knockdown of Atg16l1 or Atg5 in RAW264.7 cells significantly suppressed LRRK2 activation, not only lipidation of LC3B/GABARAP, upon apilimod treatment or the positive control of CQ treatment (**Fig. 3B and C, Supplementary Fig. 2)**. We also tested the effect of bafilomycin A_1_, which inhibits the lysosomal V-ATPase required for CASM, and confirmed its suppression of LRRK2 activation following PIKfyve inhibition (**Fig. 3D and E**). To further validate the involvement of CASM and not autophagy in LRRK2 activation, we utilized bone marrow-derived macrophages (BMDMs) from mice that express mutant Atg16l1 lacking the WD40 domain due to a stop codon inserted after E230 residue (E230/ΔWD mice) (Rai et al., 2019); these cells retain macroautophagy activity but lack CASM. We found that apilimod-induced activation of LRRK2 was suppressed in E230/ΔWD BMDMs compared to wild-type cells (**Fig. 3F and G**). Thus, these data indicate that CASM machinery is required for the activation of LRRK2 induced by PIKfyve inhibition.

We additionally found that treating cells with not only CQ but also apilimod upregulated lipidation of LC3B and GABARAP, although that of GABARAP was faint (**Fig. 3B and F, Supplementary Fig. 2)**. This lipidation was largely suppressed in E230/ΔWD BMDMs (**Fig. 3F**), indicating that PIKfyve inhibition upregulated CASM, not just the blockage of autophagy flux following lysosome dysfunction. Given that PI3K inhibition suppresses CASM-induced activation of LRRK2 but not CASM itself (**Fig. 2A-G**) and that the independency of PI3K from CASM has been confirmed (Fletcher et al., 2018; Florey et al., 2015), these data suggest that PIKfyve inhibition induces LRRK2 hyperactivation through both PI3P increase and CASM induction.

Although PI3P is not involved in typical CASM, a specific type of CASM called LC3-associated phagocytosis (LAP) is elicited through NADPH oxidase 2 (Nox2), which is activated by PI3P (Durgan and Florey, 2022; Ueyama et al., 2011). Nox2 produces reactive oxygen species following uptake of external pathogens, which then induces LAP (Huang et al., 2009) (**Fig. 3A**). Thus, it was still possible that increased PI3P could activate LRRK2 by upregulating Nox2 and then LAP. To test this possibility, we treated RAW264.7 cells with a Nox2 inhibitor diphenyleneiodonium (DPI) and examined its effect on LRRK2 activation upon PIKfyve inhibition. We found that Nox2 inhibition did not suppress apilimod-induced activation of LRRK2 (**Fig. 3H and I**), whereas it suppressed LRRK2 activation and LC3B lipidation upon treatment with zymosan, a yeast cell wall preparation that induces LAP (**Fig. 3H, J and K)**. Thus, while CASM is involved in LRRK2 activation upon PIKfyve inhibition, this does not appear to be mediated by the PI3P-Nox2 pathway upstream of LAP.

### Rab12 and Rab29 are also required for the activation of LRRK2 upon PIKfyve inhibition

It has been shown that not only CASM but also specific Rab GTPases, i.e., Rab12 and Rab32 family members (Rab29, Rab32, Rab38), are involved in LRRK2 activation. The functionally dominant Rab32 family member varies depending on the cell type, condition, etc., with Rab29 being one of essential regulators in our RAW264.7 cells (Eguchi et al., 2018; Kuwahara et al., 2020). We then examined the involvement of Rab12 and Rab29 in LRRK2 activation upon PIKfyve inhibition. As a result, knockdown of Rab12 or Rab29 in RAW264.7 cells significantly suppressed LRRK2 activation upon treatment with apilimod or YM203616 (**Fig. 4A-D**). Thus, combining the results obtained thus far, it has become clear that these Rabs, CASM and PI3P are all essential factors for the activation of LRRK2.

**Figure 4.**
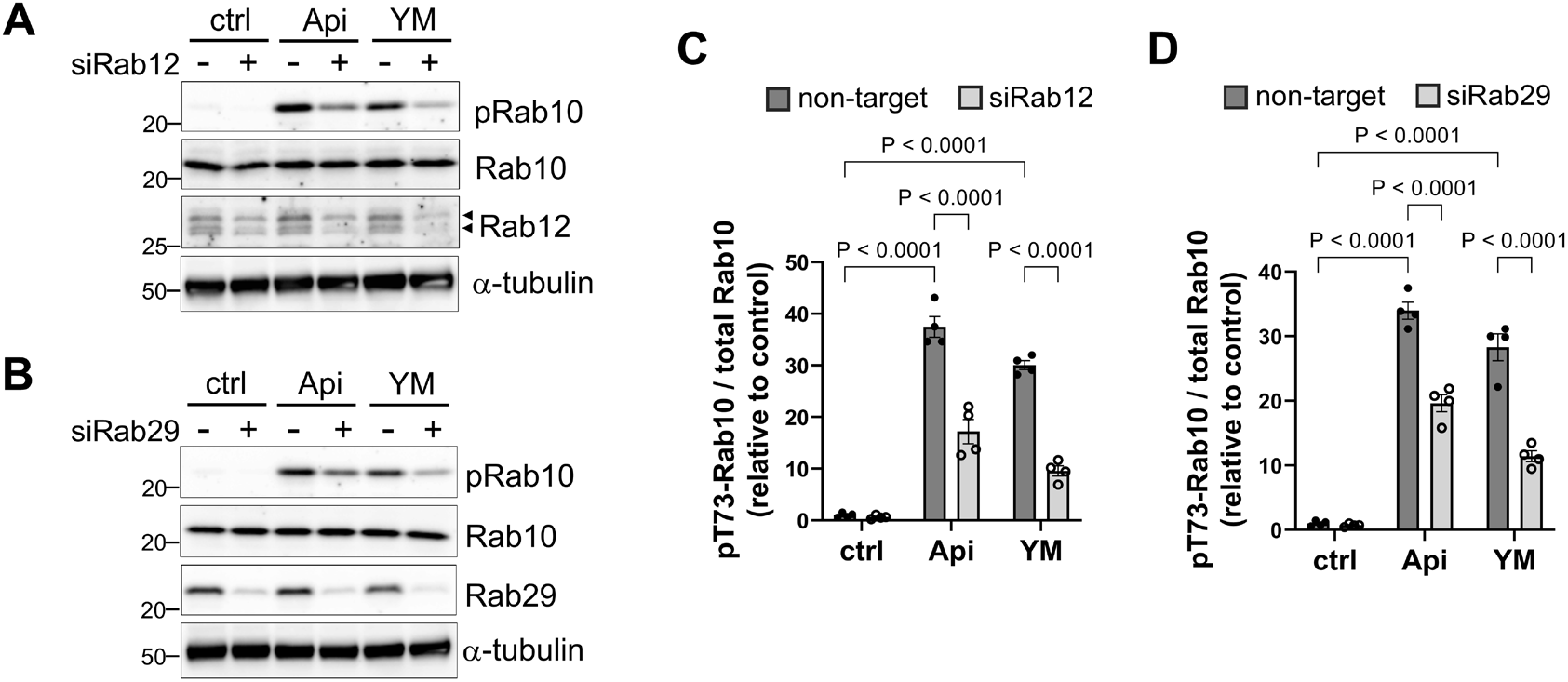
Rab12 and Rab29 are involved in LRRK2 activation upon PIKfyve inhibition. (A, **B)** Immunoblot analysis of pRab10 and other indicated proteins in RAW264.7 cells pretreated with siRNAs for Rab12 (A) or siRab29 (B) and then treated with apilimod or YM201636. **(C, D)** Quantification of pRab10 relative to total Rab10 in cells pretreated with siRab12 (D) or siRab29 (D), as shown in A and B, respectively. Data represent mean ± SEM, n = 4, two-way ANOVA with Tukey’s test.

### Endogenous LRRK2 and phospho-Rab10 colocalize with PI3P in cells exposed to PI3P-increasing stimuli

Next, we investigated whether LRRK2 is localized on PI3P-positive membranes in cells, particularly under PIKfyve inhibition. The subcellular localization of PI3P can be assessed by detecting the fluorescence of a PI3P probe p40PX-EGFP, where EGFP is fused to the PX domain of p40phox that specifically binds to PI3P (Kanai et al., 2001). We first employed HEK293 cells that are fully competent for overexpression and observed the localization of overexpressed p40PX-EGFP and 3×FLAG-LRRK2. However, we could not see their colocalization likely due to strong expression of both proteins throughout the cytoplasm, while endogenous LRRK2 in HEK293 cells was hardly detected by immunocytochemistry. Then, we examined the localization of endogenous LRRK2 in RAW264.7 cells. Since this cell line is hardly transfectable, we sought to express p40PX-EGFP by lentiviral infection, albeit at low efficiency. In this case as well, cells strongly expressing p40PX-EGFP were unsuitable for analysis, but in cells with low expression levels (where the majority of the cytoplasm had little fluorescence), endogenous LRRK2 was well colocalized with p40PX-EGFP on LAMP1-positive vacuoles formed by treatment with apilimod, YM201636 or CQ (**Fig. 5A**). This endogenous LRRK2 was not detected in *Lrrk2* knockout RAW264.7 cells, including its diffuse localization in the cytoplasm, ensuring the specificity of immunostaining **(Supplementary Fig. 3)**. We also examined the localization of endogenous Rab10 phosphorylated at Thr73, and it was similarly colocalized with p40PX-EGFP (**Fig. 5B**). Since not all signals were well matched with p40PX-EGFP, the possibility remains that other PI3P-independent mechanisms for LRRK2 recruitment exist. Nonetheless, quantitative analysis with Pearson’s correlation coefficient revealed that the colocalization of p40PX-EGFP with LRRK2 or phospho-Rab10 was increased upon treatment with apilimod, YM201636 or CQ compared to untreated controls (**Fig. 5C and D**). These colocalized proteins were found on a subset of LAMP1-positive endolysosomal vacuoles distributed throughout the cytoplasm, and the LAMP1 positivity in p40PX-EGFP-positive area was increased upon treatment with PIKfyve inhibitors or CQ (**Fig. 5E**). These observations suggest that LRRK2 becomes concentrated and activated on PI3P-rich endolysosomal membranes following PIKfyve inhibition or lysosomal stress.

**Figure 5.**
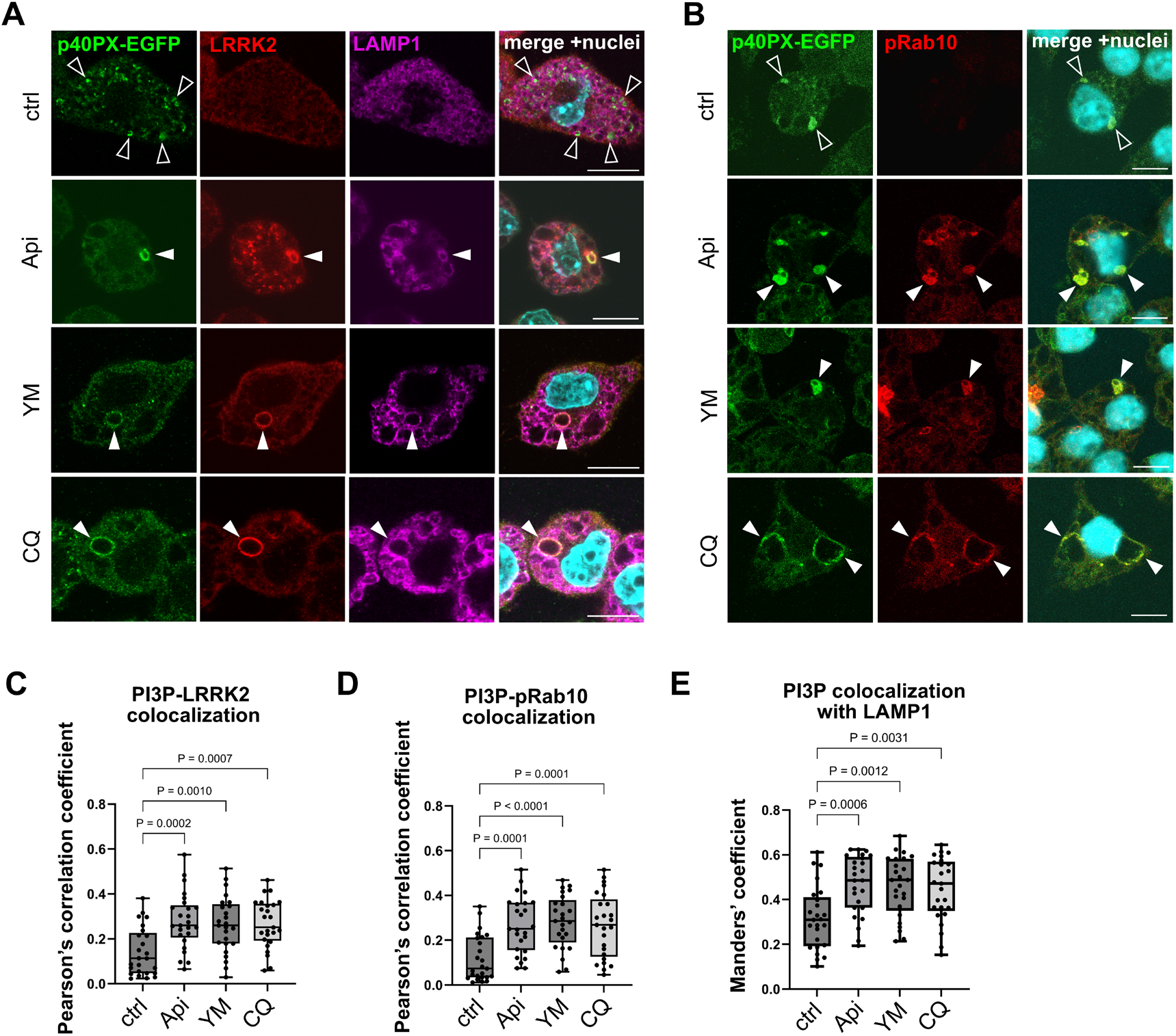
Endogenous LRRK2 and phospho-Rab10 colocalize with PI3P in cells exposed to PI3P-increasing stimuli. **(A)** Confocal microscopic analysis of the colocalization of endogenous LRRK2 (red), p40PX-EGFP (indicating PI3P, green) and LAMP1 (magenta) in RAW264.7 cells treated with PIKfyve inhibitors or CQ. Nuclei were stained with DAPI (cyan). Closed filled arrowheads indicate triple-positive vesicular structures, and closed blank arrowheads indicate PI3P-positive but LRRK2-negative structures. **(B)** Colocalization of Thr73-phosphorylated Rab10 (pRab10, red) and p40PX-EGFP (PI3P, green) in RAW264.7 cells treated with PIKfyve inhibitors or CQ. Nuclei were stained with DAPI (cyan). Closed filled arrowheads indicate pRab10-PI3P double-positive vesicular structures, and closed blank arrowheads indicate PI3P-positive but pRab10-negative structures. Scale bars in A and B = 10 μm. **(C-E)** Box-and-whisker plots indicating the colocalization of PI3P and LRRK2 (C), PI3P and pRab10 (D) or PI3P and LAMP1 (E), as shown in A and B. Colocalization shown in C and D was calculated by Pearson’s correlation coefficient for their respective fluorescent intensities, while that in E was calculated by Manders’ coefficient as the ratio of double-positive area within PI3P-positive area. n = 25 cells per group from two independent experiments, one-way ANOVA with Dunnett’s test.

### LRRK2 directly binds to PI3P via its N-terminal positively charged cluster

Since LRRK2 and PI3P were found to colocalize on stressed endolysosomes, we investigated the possibility that LRRK2 may directly bind PI3P. Recombinant human full-length LRRK2 protein with FLAG tag was reacted with nitrocellulose membranes spotted with 15 different phospholipids (each 100 pmol) including PI3P and other phosphoinositides. We found that LRRK2 reacted strongly with three phosphatidylinositol monophosphates (PI3P, PI4P, PI5P) and phosphatidic acid, with the highest reactivity on PI3P (**Fig. 6A**). This binding profile suggested that electrostatic interactions between LRRK2 and anionic lipids carrying a negative charge equivalent to one phosphate group may be involved in binding. Also, recombinant LRRK2 lacking its N-terminal 969 amino acids did not exhibit binding to the lipids on the membrane strip (**Fig. 6A**), suggesting the importance of the N-terminal domains for lipid binding. We additionally examined the binding of recombinant full-length LRRK2 with biotin-conjugated liposomes containing either PI, PI3P or PI(3,5)P_2_ by a pulldown assay. LRRK2 was specifically pulled down with liposomes containing PI3P, and, to a slightly lesser extent, with those containing PI(3,5)P_2_ (**Fig. 6B and C**).

**Figure 6.**
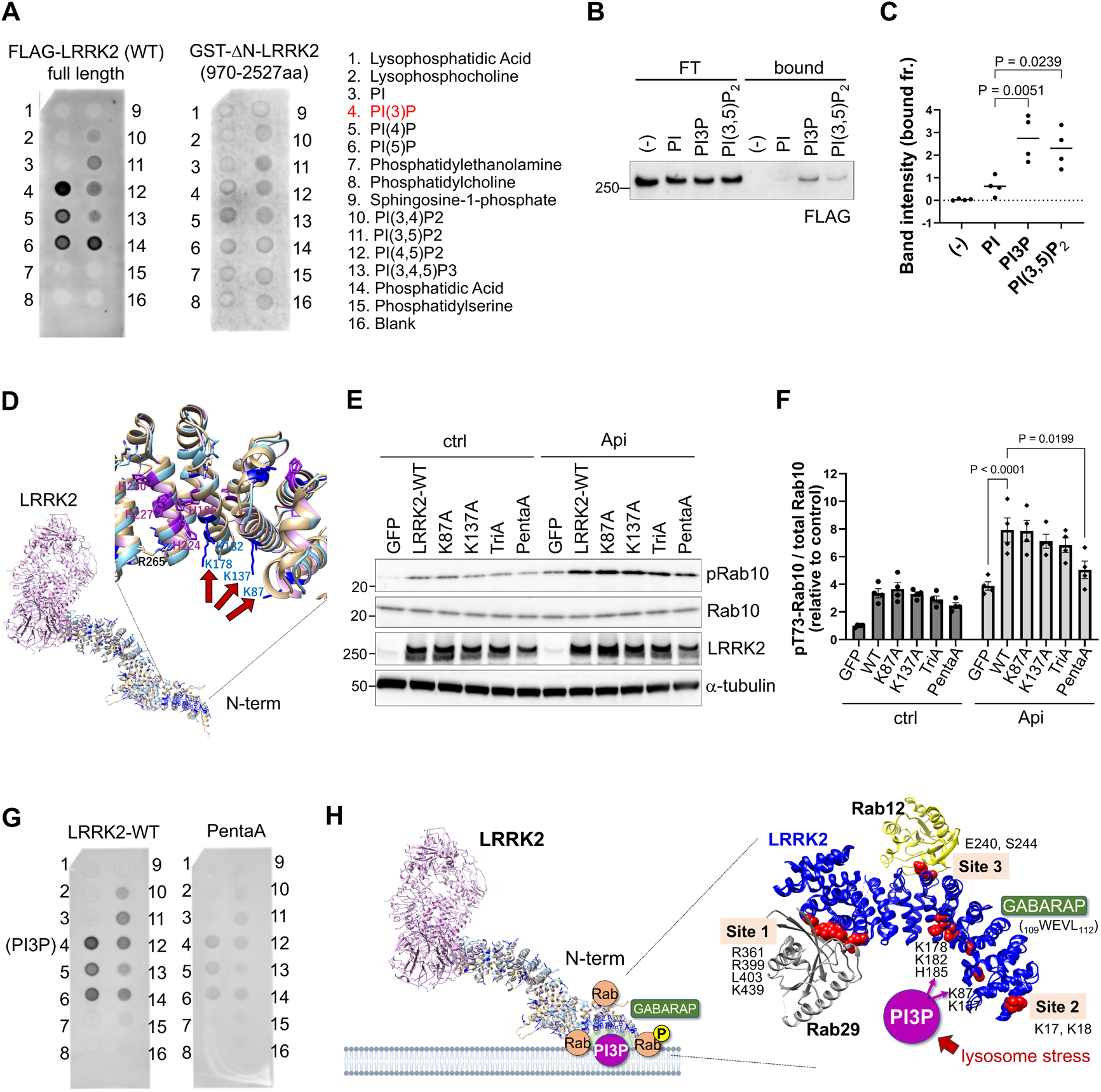
LRRK2 directly binds to PI3P via its N-terminal positively charged cluster. **(A)** Protein-lipid overlay assays for the detection of the interaction of full-length (left) or N-terminally truncated (970-2527aa, right) LRRK2 with various phosphoinositides. The names of phospholipids corresponding to each spot are shown on the right. Representative pictures from three independent experiments are shown. **(B, C)** A liposome pulldown assay for the detection of the interaction of full-length LRRK2 with biotin-conjugated liposomes containing PI, PI3P or PI(3,5)P_2_. LRRK2 bands in the flow-through (FT) and bound fractions are shown. A dot plot in C shows quantification of LRRK2 in the bound fraction, as shown in B. n = 4, one-way ANOVA with Tukey’s test. **(D)** Structure of the positively charged cluster in the N-terminal region of LRRK2, predicted to interact with PI3P. Side chains of candidate residues predicted to face the membrane are indicated by red arrows. **(E)** Analysis of apilimod-induced activation of mutant LRRK2 harboring alanine substitutions for the N-terminal positively charged amino acids in HEK293 cells. **(F)** Quantitative analysis of Thr73-phosphorylated Rab10 relative to total Rab10, as shown in C. Data represent mean ± SEM, n = 4, two-way ANOVA with Tukey’s test. **(G)** Protein-lipid overlay assays for the detection of interaction of wild-type LRRK2 (left) or PentaA LRRK2 (right) with phosphoinositides. The arrangement of phospholipids is the same as that shown in A. Representative pictures from three independent experiments are shown. **(H)** Model of LRRK2 activation via PI3P, multiple Rabs and CASM effector GABARAP on membranes. N-terminal region of LRRK2 functions as a hot spot of interaction with these molecules, mediating LRRK2 activation.

The N-terminal region of LRRK2 contains ARM and ANK repeats, and particularly the most N-terminal region in ARM repeats (12-705 aa) has been shown to interact with multiple Rab and GABARAP proteins anchored on the membrane. Structural analysis of full-length LRRK2 by cryo-electron microscopy has also predicted that the most N-terminal region faces the membrane (Myasnikov et al., 2021). However, this region does not appear to contain known PI3P/phosphoinositide-binding domains such as the FYVE domain or the Phox homology (PX) domain. We then searched for positively charged regions within the N-terminal domain that might be involved in binding to anionic lipids. Although most regions were found to be negatively charged, we confirmed the presence of clusters bearing positive charge **(Supplementary Fig. 4A)**. Also, by employing computational methods to identify positively charged amino acids with potential to interact with PI3P, we identified K87, K137 and the K178/K182/H185 cluster, whose side chains are all predicted to be oriented toward the membrane (**Fig. 6D and Supplementary Fig. 4B)**. These five amino acids are conserved across a wide range of animal species, implying important functions **(Supplementary Fig. 4B)**. We expressed mutant LRRK2 with these five amino acids substituted by alanine in HEK293 cells and first screened for those whose activation was suppressed upon apilimod treatment. Activation at 80-90% of control still occurred in LRRK2 mutants with a single Ala substitution at each amino acid or the triple mutant where K178/K182/H185 were substituted with Ala (TriA), whereas the quintuple mutant with five amino acids substituted with Ala (PentaA) reduced activation by 50% (**Fig. 6E and F**). This mutant also exhibited reduced expression levels, which was thought to result from the mutation weakening LRRK2’s membrane binding and thereby causing protein destabilization. We additionally confirmed through AlphaFold 3 predictions that the N-terminal structure is unlikely to change significantly with the PentaA mutation **(Supplementary Fig. 4C)**. Then, we generated recombinant wild-type and PentaA mutant LRRK2 protein with 3×FLAG tag purified from overexpressed HEK293 cells and evaluated their binding to phosphoinositides immobilized on nitrocellulose membranes. PentaA LRRK2 was found to exhibit an overall reduction in binding affinity for phosphoinositides, including PI3P, compared to wild-type LRRK2 (**Fig. 6G**). Thus, the five positively charged amino acids modified in PentaA appear to be involved in the binding to these phosphoinositides.

Lastly, we examined the possibility that LRRK2 behaves as a PI3P-dependent kinase in vitro, like known phosphoinositide-dependent kinases such as 3-phosphoinositide dependent protein kinase 1 (PDPK1) (Alessi et al., 1997). An in vitro kinase assay using recombinant full-length LRRK2 revealed that LRRK2 autophosphorylation activity was not changed upon addition of liposomes containing PI, PI3P or PI(3,5)P_2_ **(Supplementary Fig. 5A)**. Also, in a cell-free assay where *Lrrk2* knockout RAW264.7 cell-derived membranes were incubated with recombinant LRRK2, no changes in LRRK2 kinase activity were observed upon PI3P addition **(Supplementary Fig. 5B)**. These results indicate that the simple addition of PI3P in vitro is not sufficient to activate LRRK2 as observed in cells. Given that PI3P, CASM components and multiple Rab GTPases are properly positioned at different sites in the N-terminal region of LRRK2, a model of LRRK2 hyperactivation emerges where these essential activators bind LRRK2 simultaneously in cells to stabilize it on the endolysosomal membranes (**Fig. 6H**).

## Discussion

In this study, we revealed that LRRK2 becomes activated in cells under conditions that increase PI3P, triggered by modulation of the activities of PIKfyve and the class III PI3K. Activation of LRRK2 has previously been shown to be elicited via CASM machinery and multiple Rab GTPases, and now we added PI3P, another mediator that acts in parallel to CASM, yet both functions as essential factors. Unlike macroautophagy, CASM does not require PI3P generation; however, each component separately participates in LRRK2 activation, forming a rather complicated LRRK2 activation mechanism. In our previous study reporting CASM-dependent LRRK2 activation (Eguchi et al., 2024), we tried to show its independence from PI3K activity but unexpectedly found it to be necessary; our present finding has now solved this mystery.

It is well known that LRRK2 is translocated to the membranes of lysosomes subjected to various forms of stress, but Rab and CASM components (*e.g.*, GABARAP) alone could not fully explain its localization specificity to stressed endolysosomes. The endolysosomal membrane is rich in PI3P and PI(3,5)P_2_, with the ratio of PI(3,5)P_2_ increasing during maturation, while stress is thought to cause de-maturation to increase PI3P levels. Therefore, simultaneous binding of LRRK2 to such membranes as well as Rab and CASM components may ensure the specificity of its localization. While PI3P increase associated with endolysosomal de-maturation likely has other primary purposes, it also contributes to LRRK2 activation possibly as another rational mechanism. Once LRRK2 is properly tethered on the membrane with the aid of PI3P, CASM components and Rab, it becomes fully stabilized and activated, boosting the lysosomal stress response via Rab phosphorylation.

Our data suggested that PI3P-rich membranes, CASM components and Rab12/Rab29 all act as essential factors for LRRK2 activation. Although their detailed relationship remains debatable, these factors are thought to act both in parallel and in tandem. First, they can simultaneously bind to different sites in the N-terminal region of LRRK2, enabling their parallel action to fully stabilize LRRK2 on membranes and activate it. Thus, depletion of any single factor significantly reduced LRRK2 activation, whereas it did not completely cancel LRRK2 activation in the presence of other factors. On the other hand, our data showed that PIKfyve inhibition caused CASM induction (Fig. 3F) and CASM in turn increased PI3P levels (Fig. 2H), supporting the existence of their tandem relationship. Thus, such complex interplay among these essential factors is thought to contribute to LRRK2 hyperactivation.

The relationship between LRRK2 and phosphoinositides has not been well understood, while it has been reported that LRRK2 directly binds to acidic lipid membranes and alters their curvature (Wang et al., 2023b), supporting the findings of this study. On the other hand, different from this study, there is a report showing that inhibition of class III PI3K selectively activates PD-associated mutant LRRK2 overexpressed in HEK293 cells (Rinaldi et al., 2023). Although the effect of PI3K inhibition on mutant LRRK2 was not tested in the present study, endogenous LRRK2 activity in RAW264.7 cells in the absence of PIKfyve inhibitors showed a tendency to increase upon PI3K inhibition. This observation suggests the potential existence of additional mechanisms not directly involving PI3P when LRRK2 activity is low. Such mechanisms include the involvement of phospholipids other than PI3P, as LRRK2 exerts its activity on the intracellular membranes as long as it can bind to other phospholipids.

Activation of LRRK2 by PIKfyve inhibition has also been reported recently (Clegg et al., 2026; Malankhanova et al., 2025). These studies reported that an increase in lysosomal osmotic pressure and volume expansion activates LRRK2, and one study suggested that this increase is mediated by the activation of a chloride/proton antiporter ClC-7, which results from an expected decrease in PI(3,5)P_2_ upon PIKfyve inhibition (Clegg et al., 2026). Our present study does not rule out the possibility of a local decrease in PI(3,5)P_2_, while our bulk quantification of PIPs showed that the increase in PI3P was the only significant change and correlated with LRRK2 activation (Fig. 2H). Nonetheless, an increase in osmotic pressure is highly plausible in our PIKfyve inhibition experiments as well, and this may accompany an increase in PI3P. Another difference between our data and the aforementioned paper is the necessity of CASM in LRRK2 activation upon PIKfyve inhibition. Our results, especially the finding that this LRRK2 activation is suppressed in ATG16L1 ΔWD40 cells that specifically lack CASM, strongly support the role of CASM as a contributing factor. This is further supported by previous findings that linked lysosomal osmotic stress to CASM (Florey et al., 2015). A commentary on the above study also stated LRRK2 activation under PIKfyve inhibition as CASM-free (Pfeffer, 2026), but this needs to be carefully verified through independent studies before it becomes established as general idea.

Although PI3P-mediated activation model is promising, another point requiring caution is whether direct binding of LRRK2 to PI3P in vitro can sufficiently explain its activation in cells. First, it remains unclear whether the binding pattern in vitro is identical to that occurring within cells. Second, even if binding occurs as expected in cells, LRRK2 activation may occur not only via this binding but also through other PI3P-dependent pathways. Nonetheless, LRRK2 PentaA mutation predicted to abolish its binding to PI3P significantly suppressed LRRK2 activation, supporting the notion that binding and activity are linked. Furthermore, we tested the possibility that PI3P might indirectly activate LRRK2 via NOX2-mediated CASM activation, but this was ruled out. Therefore, the model of LRRK2 activation through direct binding to PI3P appears to be most plausible.

We also conducted computational analysis to predict candidate N-terminal amino acids in LRRK2 that could bind to PI3P, and we indeed found evidence of their involvement in binding. Although *in silico* prediction in the absence of known PI3P-interacting motifs leaves open the possibility of additional binding sites, the predicted amino acids differ from known binding sites for Rab12, Rab29, phospho-Rab or GABARAP and are consistent with the predicted orientation toward the membrane. We should also note that, while the binding is thought to be mediated by electrostatic interactions, the positively charged binding site is located in a structurally slightly indented region, and the surrounding area has a higher negative charge. This may explain why it readily binds to monophosphates such as PI3P, which carries a weaker negative charge compared to diphosphates and triphosphates. This line of reasoning in turn suggests that LRRK2 may bind to monophosphates PI4P and PI5P as well, and indeed, this was observed in vitro. Thus, the possibility remains that LRRK2 additionally localizes to membranes rich in such lipids. For example, PI4P is known to increase in lysosomal membranes damaged by LLOMe and other lysosomal membrane-permeabilizing agents (Radulovic et al., 2022; Tan and Finkel, 2022), and LRRK2 may bind to such phosphoinositide to localize there because LRRK2 is known to be activated also by LLOMe treatment (Herbst et al., 2020; Kalogeropulou et al., 2020). This possibility is worth further consideration in the future.

The relationship between PD pathomechanisms and PI3P-associated LRRK2 activation is also of interest. Given that elevated LRRK2 kinase activity has been reported in sporadic PD tissues and cells (Di Maio et al., 2018; Petropoulou-Vathi et al., 2022), it is possible that PI3P levels are increased at the lesion site. Alternatively, the endolysosomal localization of Rabs or CASM components may be increased there. Since lysosomal dysfunction in human cells is known to progress with aging, as evidenced by the accumulation of lipofuscin granules, SA-β-gal, etc. (Goyal, 1982; Kurz et al., 2000), such dysfunction may lead to increased PI3P and/or CASM, potentially raising the risk of LRRK2-associated PD. Therefore, it is desirable to investigate these possibilities using human patients-derived samples in the future. In terms of drug discovery, compounds that block the interaction between LRRK2 and PI3P could potentially yield drugs that suppress abnormal activation of LRRK2 without inhibiting its basal activity. Such an approach may also serve as a possible direction for future research in disease studies.

## Materials and Methods

### Antibodies and reagents

The following antibodies were used in this study. The dilution ratios shown are for immunoblotting, unless otherwise specified. anti-LRRK2 [MJFF2 (c41-2)] (Abcam, 1:1,000 for immunoblotting, 1:200 for immunocytochemistry), anti-phospho-Thr73 Rab10 [MJF-R21 (ab230261)] (Abcam, 1:1,000 for immunoblotting, 1:200 for immunocytochemistry), anti-Rab10 [D36C4] (Cell Signaling, 1:1,000), anti-Rab10 [4E2 (ab104859)] (Abcam, 1:1,000), anti-phospho-Rab [anti-phospho-Thr72 Rab8A, MJF-R20] (Abcam, 1:1,000), anti-α-tubulin [DM1A] (Sigma, 1:10,000), anti-mouse LAMP1 [1D4B] (Bio-Rad, 1:3,000 for immunoblotting, 1:500 for immunocytochemistry), anti-LC3 [L7543] (sigma, 1:2000), anti-GABARAP [E1J4E] (Cell Signaling, 1:500), anti-ATG16L1 [D6D5] (Cell Signaling, 1:1000), anti-ATG5 [A0731] (Sigma, 1:1000), anti-Rab12 [18843-1-AP] (Proteintech, 1:500), anti-Rab29 [MJF-R30-124] (Abcam, 1:1000), anti-FLAG [M2] (Sigma), anti-GST [27-4577-01] (Cytiva, 1:5000), anti-phospho-Thr [#9381] (Cell Signaling, 1:1000), anti-pS1292 LRRK2 [ab203181] (Abcam, 1:500). The following reagents were used at final concentrations as indicated: apilimod (100 nM, MedChemExpress), YM203616 (5 μM, AdooQ), VPS34-IN1 (10 μM, Cayman), Compound 19 (50 μM, Selleck), 3-methyladenine (3 mM, AdipoGen), chloroquine (50-100 μM, Sigma), monensin (50 μM, Cayman), nigericin (10 μM, Sigma), diphenyleneiodonium chloride (DPI, 10 μM, Selleck), zymosan (Sigma), bafilomycin A_1_ (30 nM, Wako).

### Plasmids and siRNAs

Plasmids encoding 3×FLAG-human LRRK2 WT, R1441C and Y1699C were used previously (Fujimoto et al., 2018; Kuwahara et al., 2020). To construct plasmids encoding 3×FLAG-human LRRK2 with an alanine substitution mutation at the N-terminal region, p3×FLAG-human LRRK2 WT were first cut by Bsp1407 I (Takara Bio) to remove LRRK2 N-terminal region, and then the two PCR amplicons overlapping the mutation site as well as the cleaved plasmid were assembled using NEBuilder (New England Biolabs). To construct a lentiviral plasmid for the expression of p40PX-EGFP in RAW264.7 cells, p40PX-EGFP sequence (Addgene) was PCR-amplified and inserted into the Xho I-Not I site of the pLVSIN-CMV Neo vector (Takara Bio) using Ligation high Ver.2 (Toyobo). All constructs were sequence-verified. siRNAs for the knockdown of gene expression were purchased from Horizon Discovery (Dharmacon siGENOME, *Pikfyve*: Cat#D-040127-05 and D-040127-06, *Atg5*: Cat#D-064838-01, *Atg16l1*: Cat#D-051699-01, *Rab12*: Cat#M-040865-01, *Rab29*: Cat#M-053101-00).

### Cell culture and transfection

RAW264.7 mouse macrophage cells (ECACC, RRID: CVCL_0493) and *Lrrk2* knockout RAW264.7 cells (ATCC, RRID: CVCL_UL72) were routinely cultured on culture dishes for suspended cells (Sumitomo Bakelite Co.) in Dulbecco’s modified Eagle’s medium (DMEM) supplemented with 10% (v/v) fetal bovine serum (FBS). These cells were pretreated with interferon-γ (15 ng/ml, Cell Signaling) for 24-48 h prior to assay, which was necessary to fully elicit the activity of endogenous LRRK2. Human embryonic kidney (HEK) 293 cells (ATCC, RRID: CVCL_0045) and human lung carcinoma A549 cells (ECACC, RRID: CVCL_0023) were cultured on normal dishes in DMEM supplemented with 10% (v/v) FBS. Bone marrow-derived macrophages (BMDMs) were generated from femurs and tibias of wile-type and E230 mice as described previously (Eguchi et al., 2024). Briefly, macrophages were generated from adherent cells in RPMI-1640 containing 10% FCS and M-CSF (Peprotech, 315-02) (30 ng/ml) for 6 d. Macrophage populations were quantified by FACS using antibodies against CD16/CD32, F4/80 and CD11b (BioLegend, 101320, 123107). The experiment of BMDMs isolation from mice was performed in accordance with UK Home Office guidelines and under the UK Animals (Scientific procedures) Act1986. All cell lines were cultured at 37°C in a 5% CO_2_ atmosphere. Chemical reagents were treated to cells for 2-3 h by medium exchange. Transfection of LRRK2 plasmids in HEK293 cells was performed using Lipofectamine 3000 (Thermo Fisher), according to the manufacturer’s protocols, and cells were analyzed after 24-48 h. Transfection of siRNAs was performed using Lipofectamine RNAiMAX (Thermo Fisher), and cells were analyzed after 48-72 h. For immunocytochemistry, cells were seeded on a coverslip without any special coating.

### Lentiviral expression of p40PX-EGFP

For the production of recombinant lentiviral particles, pLVSIN-CMV Neo vector encoding p40PX-EGFP and Lentivial High Titer Packaging Mix (Takara Bio) were transfected into Lenti-X 293T cells (Takara Bio) using Lipofectamine 3000 (Thermo Fisher), according to the manufacturer’s protocols. Medium was changed after 6-12 h of transfection, then medium was collected and refreshed daily for 2 d. The lentiviral particles in medium were concentrated using Lenti-X Concentrator (Takara Bio), resuspended in DMEM, and then immediately infected to RAW264.7 cells together with polybrene (Millipore). The medium was changed after 24 h, and the infected cells were analyzed the following day.

### SDS-PAGE and immunoblot analysis

Immunoblot analysis basically follows our previous method (Abe et al., 2024; Sakurai and Kuwahara, 2021). Cells were washed with PBS on ice and lysed in a lysis buffer containing 50 mM Tris HCl pH 7.6, 150 mM NaCl, 1% (v/v) Triton X-100, Complete EDTA-free protease inhibitor cocktail (Roche), and PhosSTOP phosphatase inhibitor Cocktail (Roche). Lysates were centrifuged at 17,800 × *g* for 5 min at 4°C and supernatants were mixed with NuPAGE LDS sample buffer (4×) (Thermo Fishe). For SDS-PAGE, samples were loaded onto polyacrylamide gels (SuperSep Ace, Fujifilm Wako) and electrophoresed. After electrophoresis, samples were transferred to PVDF membranes using iBlot3 (Invitrogen). Transferred membranes were blocked with 5% skim milk (BD), incubated with primary antibodies and then with horseradish peroxidase (HRP)-conjugated secondary antibodies (Jackson ImmunoResearch). Membranes were incubated with ImmunoStar Zeta (Fujifilm Wako) and the chemiluminescence of the protein bands was detected using LAS-4000 (Fujifilm). For densitometric analysis, the integrated densities of protein bands were calculated using ImageJ software (NIH). To further align the overall chemiluminescence intensity among several different membranes, the band intensities were normalized so that the sum of the intensities of all bands for each membrane was equal.

### Measurement of cellular phosphoinositide levels by mass spectrometry

RAW264.7 cells cultured on two 10 cm-diameter culture dishes (for suspended cells, Sumitomo Bakelite Co.) were treated with each drug for 3 h, then cells were harvested using a cell scraper. After counting the cells, they were centrifuged at 200 × *g* for 3 min, the supernatant was discarded, and the cell pellet was temporally stored at -80°C. The subsequent analysis was conducted at Lipidome Lab Co., Ltd, as described previously (Morioka et al., 2022). Briefly, samples were dissolved in methanol, and total lipids were extracted by the modified Bligh and Dyer method. The total lipid fraction was further purified and recovered using an anion exchange column to isolate the acidic phospholipid fraction including phosphoinositides. The purified fraction was dried under a stream of nitrogen, then redissolved in acetonitrile and transferred to vials. LC-MS/MS analysis was performed using the Xevo TQ-XS mass spectrometer with an ACQUITY UPLC H-Class (Waters). The lipids were separated using a CHIRALPAK IC-3 column (2.1 × 250 mm, 3 μm, DAICEL) at 23°C under the conditions as described (Morioka et al., 2022). Phosphoinositides species were measured using multiple reaction monitoring (MRM) in positive ion mode, and peak picking was conducted using analytical software MassLynx4.2 (Waters). Peak areas of individual species were normalized with those of the internal/surrogate standards, and final quantification data were obtained by correcting for the total phosphatidylserine (PS) amount detected and presented in pmol / PS 1000 pmol units.

### Immunocytochemistry

Cells cultured on coverslips were fixed with 4% (w/v) paraformaldehyde for 20 min, followed by immersion in 100% EtOH at -20°C, which was necessary for the staining with an anti-LRRK2 antibody c41-2 [MJFF2]. Samples were washed with PBS and then permeabilized and blocked with 3% (w/v) BSA in PBS containing 0.1% Triton X-100. Samples were incubated with primary antibody solutions diluted at the ratio as described above, followed by incubation with corresponding secondary antibodies conjugated with fluorescent dyes (goat anti-rabbit IgG Alexa Fluor 546 and goat anti-rat IgG Alexa Fluor 647, Thermo Fisher) diluted at 1:1000 in the blocking buffer. Nuclei were stained with DAPI (Thermo Fisher) at 1:5000 dilution. The samples were imaged using a confocal laser scanning microscope (FLUOVIEW FV3000, Evident) equipped with a 100× oil immersion objective lens. Image contrast and brightness were adjusted using Photoshop 2025 software (Adobe). Colocalization of p40PX-EGFP fluorescence and endogenous LRRK2/pRab10 signal was measured using Pearson’s correlation coefficient, while colocalization of p40PX-EGFP and LAMP1 was measured using Manders’ coefficient, both using the ImageJ (NIH) JACoP plugin with Costes’ automatic threshold method.

### Purification of LRRK2 protein

Human LRRK2 WT and PentaA proteins tagged with 3×FLAG at N-terminus were purified from monoclonal HEK293 cells stably expressing either 3×FLAG-LRRK2 WT or PentaA as follows. First, these cells cultured on ten 15 cm diameter dishes were washed once with D-PBS, collected into 10 ml (per dish) of D-PBS using cell scraper (SPL) and centrifuged at 200 × *g* for 3 min. Cell pellets were then lysed in 12.5 ml of ice-cold lysis buffer (TBS (50 mM Tris-HCl pH 7.6, 150 mM NaCl), 0.5 mM EDTA, 0.5% (v/v) Triton X-100, Complete EDTA-free protease inhibitor cocktail (Roche), PhosSTOP phosphatase inhibitor cocktail (Roche), 1 mM PMSF (Wako)) and rotated for 1 h at 4°C. The lysates were centrifuged at 17,800 × *g* for 5 min at 4°C, and after collecting a small aliquot as the input fraction, supernatants were mixed with 100 ul of TBS-equilibrated anti-FLAG M2 Affinity Gel (Merck-Millipore) and rotated overnight at 4°C. Samples were then centrifuged at 5,800 × *g* for 1 min at 4°C, and the gel pellets were resuspended in 1.6 ml of ice-cold wash buffer (TBS (50 mM Tris-HCl pH 7.6, 150 mM NaCl), 0.5 mM EDTA, 0.1% (v/v) Triton X-100, 1 mM PMSF (Wako)). After repeating this wash step four times, the gel pellets were mixed with 200 ul of 3×FLAG peptide (200 μg/ml in TBS, Sigma) by pipetting and incubated for 20 min at room temperature with rotation. Samples were centrifuged at 5,800 × *g* for 1 min at 4°C, and the supernatant was transferred to new tubes. The gel pellets were again mixed with 160 ul of 3×FLAG peptide, and the procedure of incubation followed by centrifugation was repeated to obtain the supernatant from two runs as the elute fraction. To remove 3×FLAG peptide carried over, this fraction was further added to Nanosep 30K Omega centrifugal filters (Pall Life Sciences) and centrifuged at 2000 × *g* for 20 min at 4°C. Samples remaining in the upper chamber were mixed with 450 ul of TBS and centrifuged again, and after repeating this step twice, the samples in the upper chamber were recovered as the purified concentrated fraction.

### Protein-lipid overlay assay

The direct binding of LRRK2 to various phospholipids was assessed using PIP Strips (P-6001, Echelon Biosciences). PIP membranes were first blocked with blocking buffer containing 3% bovine serum albumin (BSA) in TBS-T (0.1% (v/v) Tween 20 in TBS) for 1 h and then probed with 1 μg of recombinant FLAG-tagged full-length LRRK2 WT (Invitrogen, and the protein purified in house for comparison with PentaA), LRRK2 PentaA (purified in house) or GST-tagged ΔN-LRRK2 (970-2527 aa) (Invitrogen) in blocking buffer for 1 h. Membranes were then washed with TBS-T three times and incubated with anti-FLAG antibody (Sigma) or anti-GST antibody (Cytiva) for 1 h. After washing with TBS-T three times, membranes were incubated with HRP-conjugated secondary antibody (Jackson ImmunoResearch) for 1 h and then washed again with TBS-T three times. Membranes were incubated with ImmunoStar Zeta (Fujifilm Wako) and the chemiluminescence of the signal was detected using LAS-4000 (Fujifilm).

### Liposome pulldown assay

Recombinant FLAG-tagged full-length LRRK2 WT (0.25 μg, Invitrogen) was mixed with or without 7.5 ul of biotin-conjugated polymerized liposomes containing PI, PI3P or PI(3,5)P_2_ (PolyPIPosome, Echelon Biosciences) in 300 μl of the binding buffer (0.05% (v/v) NP-40 in TBS). After 15 min of incubation with rotation at room temperature, the samples were mixed with 15 ul of streptavidin-coupled magnetic beads (Dynabeads MyOne Streptavidin T1, Thermo Fisher) prewashed twice with TBS and then incubated for 30 min with rotation at room temperature. The samples were washed four times with the binding buffer using a magnetic rack, while the supernatant after the first wash was collected as the flow-through fraction. After removing buffer, the samples with beads were resuspended in 25 ul of NuPAGE LDS sample buffer (Thermo Fisher) and incubated for 5 min at 90°C. The samples were then placed on a magnetic rack for 1-2 min, and the supernatants were collected as the bound fraction. The presence of FLAG-LRRK2 in the flow-through and the bound fractions were analyzed by SDS-PAGE and immunoblotting using an anti-FLAG M2 antibody.

### *In silico* prediction of the LRRK2-PI3P interaction site

The DiffDock-L (Corso et al., 2024), deep-learning-based docking method, was used to predict the interaction sites of LRRK2 with PI3P using the N-terminal portion of 325 residues from the crystal structure of LRRK2 (C chain of 8FO8) and Ins(1,3)P_2_, a soluble mimic of PI3P, for the prediction. Note that the crystal structure of LRRK2 lacks a few lysine residues on its α-helix. The prediction was performed using DiffDock-Web on Hugging Face Spaces with the default parameters. Candidate sites were narrowed down by taking into account the conservation of residues among homologous proteins in vertebrates. The multiple sequence alignment of homologous proteins was calculated with FAMSA (Deorowicz et al., 2016). The surface charges of LRRK2 were calculated and visualized using UCSF Chimera.

### In vitro/cell-free kinase assay

For in vitro kinase assay evaluating LRRK2 autophosphorylation, recombinant FLAG-LRRK2 (200 ng, Invitrogen) was mixed with 4 μg of liposomes containing 5% PI, PI3P or PI(3,5)P_2_ (PolyPIPosome, Echelon Biosciences) in 20 μl of an assay buffer (50 mM Tris-HCl pH7.5, 10 mM MgCl_2_, 1 mM CaCl_2_, 2 mM DTT, 1mM ATP) and incubated for 20 min at 30°C while shaking at 300 rpm using Thermo Shaker BSR-MS100 (Biomedical Science). Samples were then subjected to immunoblot analysis using anti-phospho-Thr and anti-FLAG antibodies. For cell free kinase assays, membrane fractions from *Lrrk2* knockout RAW264.7 cells were first prepared by homogenizing cell pellets in a homogenization buffer (20mM HEPES-KOH (pH7.5), 250 mM sucrose, cOmplete EDTA-free protease inhibitor cocktail (Roche)) followed by centrifugation at 800 × *g* for 10 min (collecting sup) and then ultracentrifugation at 100,000 × *g* for 15 min. The pellet was resuspended in a kinase assay buffer (50 mM HEPES-KOH (pH7.5), 10 mM MgCl_2_, 1 mM CaCl_2_, 2mM DTT, 1mM ATP, cOmplete EDTA-free, PhosSTOP (Roche)), and then recombinant FLAG-LRRK2 G2019S (5 nM, Invitrogen) and PI(3)P diC16 (Echelon Biosciences) at several concentrations were added to 30 μl of the mixture. Samples were incubated for 60 min at 30°C while shaking at 400 rpm and subjected to the immunoblot analysis using anti-phospho-Rab and other antibodies.

## Statistics

The statistical significance of difference in mean values was calculated by one-way or two-way ANOVA with Tukey’s post hoc test using GraphPad Prism 9. Standard error of the mean (SEM) was shown in all graphs. *P* values less than 0.05 were considered statistically significant. No exclusion criteria were applied to exclude samples from analysis.

## Data availability

All data sets used or analyzed in this study are available from the corresponding author upon request.

## Acknowledgements

The authors thank the Iwatsubo lab members as well as Prof. Takashi Saito’s lab members for helpful suggestions and discussions. This study was supported by JSPS KAKENHI grant numbers 20H00525 (T. I.), 22H02949 (T. K.), 25K02458 (T. K.), ONO Medical Research foundation (T. K.), Takeda Science Foundation (T. K.), and by Research Support Project for Life Science and Drug Discovery (Basis for Supporting Innovative Drug Discovery and Life Science Research (BINDS)) from AMED under Grant Number JP25ama121028 (support number 6300) (K. T.).

## Author contributions

Conceptualization: T. K. and G. Y.; Investigation: T. K., G. Y., M. S., S. S., H. N. and K. T.; Methodology: T. K., H. N. and K. T.; Resources: M. J., T. W.; Data curation: T. K., G. Y., H. N., K. T. and T. I.; Writing – original draft: T. K.; Writing – review & editing: T. K., H. N., T. W., K. T. and T. I.; Supervision: T. K. and T. I.; Project administration: T. K.; Funding acquisition: T. K. and T. I. All authors read and approved of the final manuscript.

