## Supplementary Figures for "PIKfyve inhibition induces LRRK2 activation through phosphatidylinositol 3-phosphate and CASM"

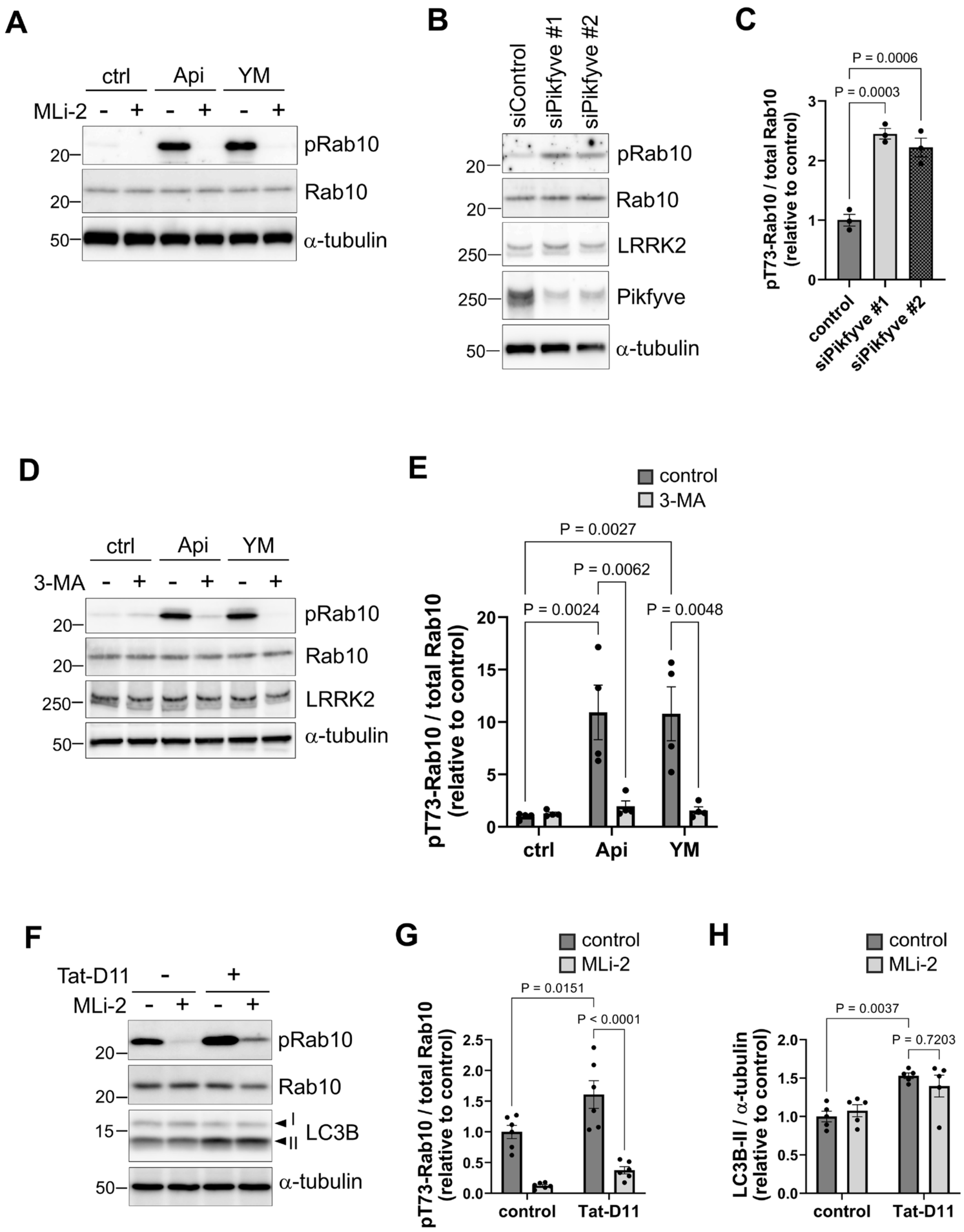


**Figure S1. Effects of MLi-2 and 3-MA on Rab10 phosphorylation upon treatment with PIKfyve inhibitors or Tat-D11.**

**(A)** Immunoblot analysis of Thr73-phosphorylated Rab10 (pRab10), Rab10 and a-tubulin in RAW264.7 cells treated with PIKfyve inhibitors and MLi-2, a LRRK2-specific inhibitor. **(B, C)** Immunoblot analysis of pRab10 and other indicated proteins in RAW264.7 cells treated with two different siRNAs for Pikfyve. A bar graph in C shows quantification of pRab10 relative to total Rab10, as shown in B. Data represent mean ± SEM, n = 3, one-way ANOVA with Dunnett's test. **(D, E)** Immunoblot analysis of pRab10 and other indicated proteins in RAW264.7 cells treated with PIKfyve inhibitors and 3-MA, a PI3K inhibitor. A bar graph in E shows quantification of pRab10 relative to total Rab10, as shown in D. Data represent mean ± SEM, n = 4, two-way ANOVA with Tukey's test. **(F-H)** Immunoblot analysis of pRab10, LC3B and other indicated proteins upon treatment with Tat-D11 and/or MLi-2 in RAW264.7 cells. Bar graphs show quantifications of pRab10 relative to total Rab10 (G) and LC3B-II relative to α-tubulin (H). Data represent mean ± SEM, n = 6 (G) or 5 (H), two-way ANOVA with Tukey's test.

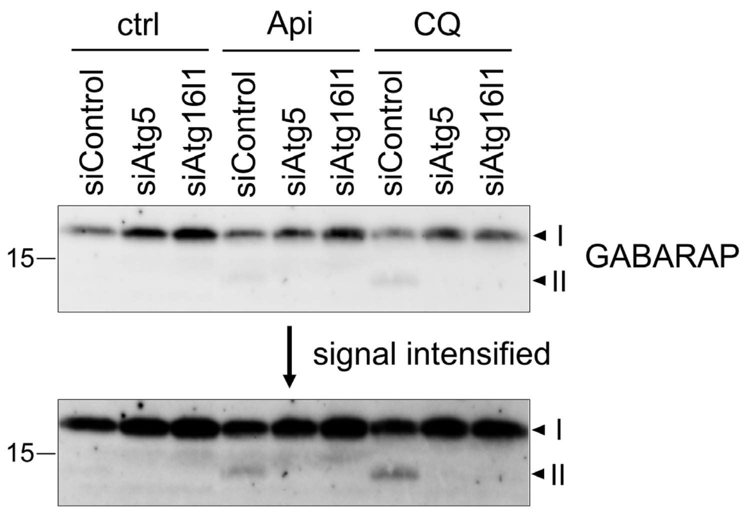


**Figure S2. Lipidation of GABARAP under treatment with siAtg5/siAtg16l1 and then with apilimod or CQ, as shown in Fig. 3B.**
Immunoblot analysis of GABARAP in RAW264.7 cells treated with the indicated siRNAs and reagents. A representative image from three independent experiments, together with a modified image with the signal intensified, is shown. α-Tubulin band as a loading control is shown in Fig. 3B.

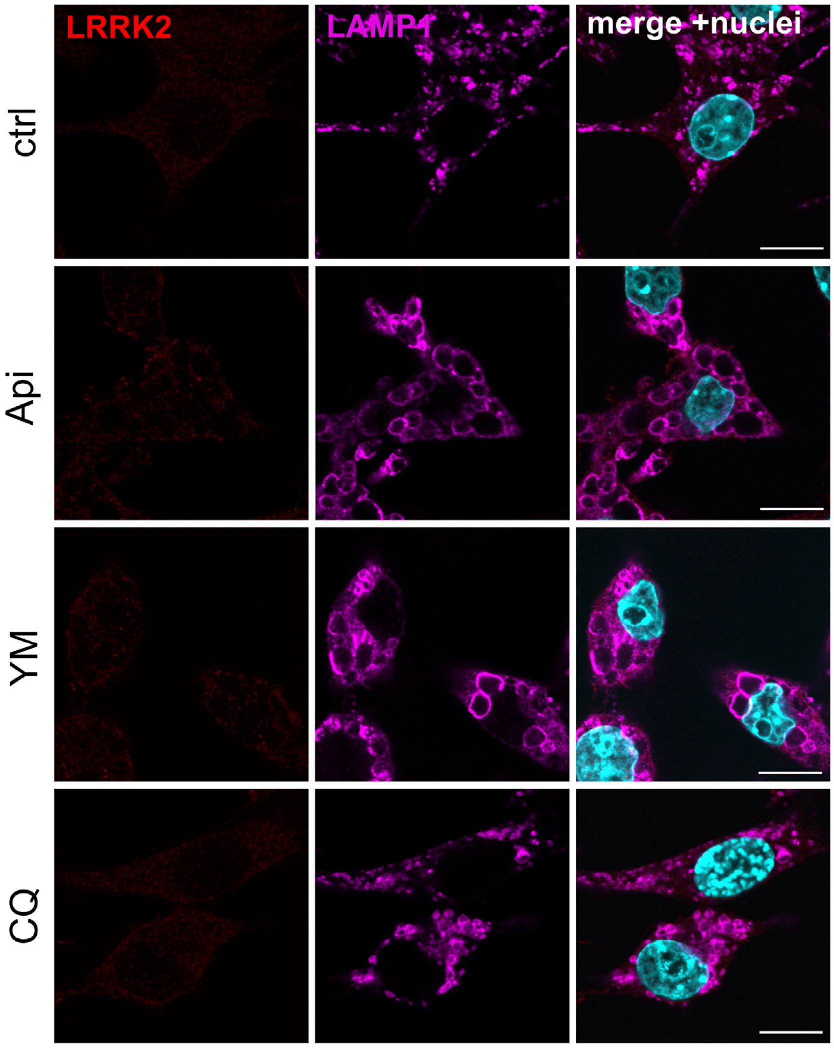


**Figure S3. Lack of LRRK2 immunoreactivity in *Lrrk2* KO RAW264.7 cells.** Confocal microscopic images of *Lrrk2* knockout (KO) RAW264.7 cells treated with PIKfyve inhibitors or CQ, fixed and then immunostained for endogenous LRRK2 (red) and LAMP1 (magenta). Nuclei were stained with DAPI (cyan). Scale bars = 10 µm.

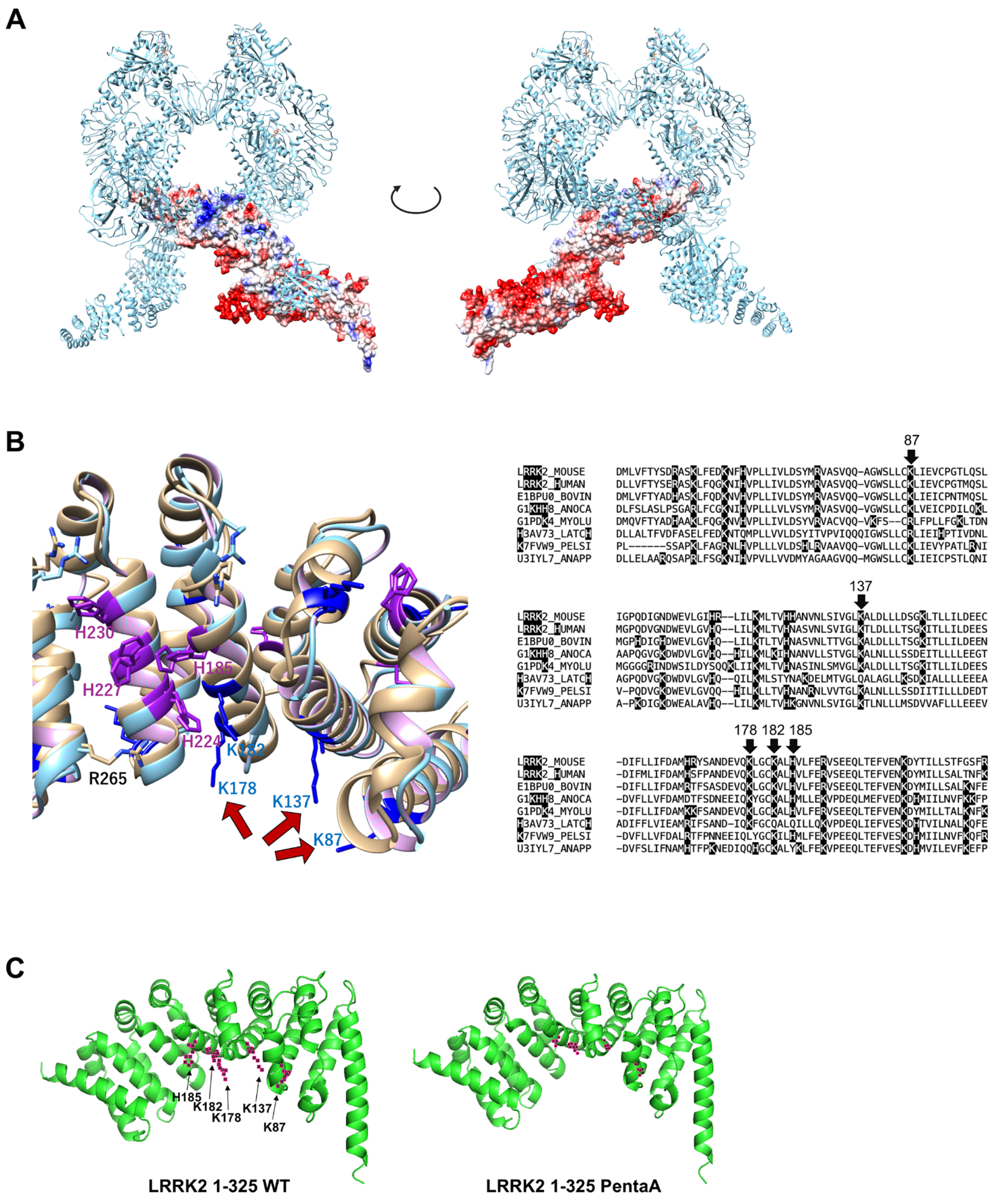


**Figure S4. In silico analysis of the binding site between LRRK2 and PI3P.**

**(A)** The charged state of full-length human LRRK2 dimer (based on PDB: 8FO8), as calculated and visualized using UCSF Chimera. Red indicates negatively charged areas, while blue indicates positively charged areas. **(B)** Left: the enlarged view of the predicted binding site for PI3P, related to Figure 6B. The side chains of candidate amino acids are indicated by red arrows. Right: multiple sequence alignments showing conservation of candidate PI3P-binding amino acids across species. **(C)** Comparison of the N-terminal structures of the wild-type and PentaA mutant LRRK2 (1-325 amino acids) predicted by AlphaFold 3.

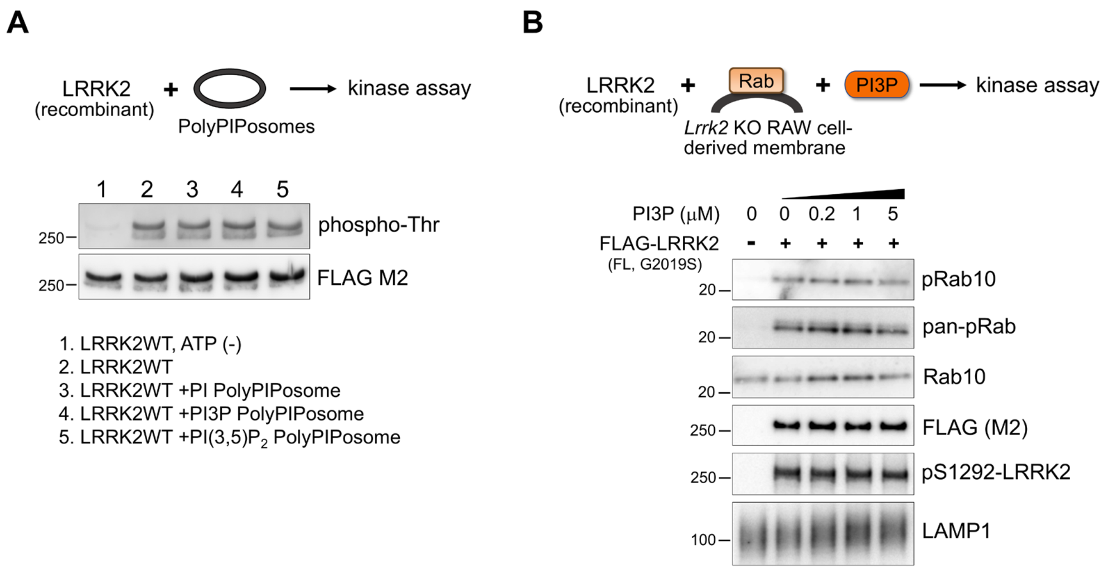


**Figure S5. LRRK2 in vitro/cell-free kinase assays.**

**(A)** An in vitro kinase assay using recombinant FLAG-LRRK2 and liposomes containing PI, PI3P or PI(3,5)$P_2$ (PolyPIPosomes). The sample without addition of ATP was included as a negative control. **(B)** A cell-free kinase assay using recombinant FLAG-LRRK2 and *Lrrk2* KO RAW264.7 cell-derived membrane fractions with various concentrations of PI3P.

**Table S1.** Full lipidomics results for the quantitative analysis of phosphoinositide regioisomers using chiral column chromatography and mass spectrometry.
